# Antisense oligonucleotides against monoacylglycerol acyltransferase 1 (*Mogat1*) improve glucose metabolism independently of *Mogat1*

**DOI:** 10.1101/2020.08.05.238535

**Authors:** Andrew J. Lutkewitte, Jason M. Singer, Trevor M. Shew, Michael R. Martino, Angela M. Hall, Brian N. Finck

**Author notes:** Acceptance of proofs and correspondence to: Brian N. Finck, Center for Human Nutrition, Washington University School of Medicine, 660 Euclid Ave., Box 8031, St. Louis, MO 63110, United States of America.

## Abstract

**Objective:** Monoacylglycerol acyltransferase (MGAT) enzymes catalyze the synthesis of diacylglycerol from monoacylglycerol. Previous work has suggested the importance of MGAT activity in the development of obesity-related hepatic insulin resistance. Indeed, antisense oligonucleotide (ASO)-mediated knockdown of the gene encoding MGAT1, *Mogat1*, reduced hepatic MGAT activity and improved glucose tolerance and insulin resistance in high fat diet (HFD) fed mice. However, recent work has suggested that some ASOs may have off-target effects on body weight and metabolic parameters via activation of the interferon alpha/beta receptor 1 (IFNAR-1) pathway.

**Methods:** Mice with whole-body *Mogat1* knockout or a floxed allele for *Mogat1* to allow for liver-specific *Mogat1*-knockout (by either a liver-specific transgenic or adeno-associated virus-driven Cre recombinase) were generated. These mice were placed on a high fat diet and glucose metabolism and insulin sensitivity was assessed after 16 weeks on diet. In some experiments, mice were treated with control or *Mogat1* or control ASOs in the presence or absence of IFNAR-1 neutralizing antibody.

**Results:** Genetic deletion of hepatic *Mogat1*, either acutely or chronically, did not improve hepatic steatosis, glucose tolerance, or insulin sensitivity in HFD-fed mice. Furthermore, constitutive *Mogat1* knockout in all tissues actually exacerbated HFD-induced weight gain, insulin resistance, and glucose intolerance on a HFD. Despite markedly reduced *Mogat1* expression, liver MGAT activity was unaffected in all knockout mouse models. *Mogat1* overexpression hepatocytes increased liver MGAT activity and TAG content in low-fat fed mice, but did not cause insulin resistance. Interestingly, *Mogat1* ASO treatment improved glucose tolerance in both wild-type and *Mogat1* null mice, suggesting an off target effect. Inhibition of IFNAR-1 did not block the effect of *Mogat1* ASO on glucose homeostasis.

**Conclusion:** These results indicate that genetic loss of *Mogat1* does not affect hepatic MGAT activity or metabolic homeostasis on HFD and show that *Mogat1* ASOs improve glucose metabolism through effects independent of targeting *Mogat1* or activation of IFNAR-1 signaling.

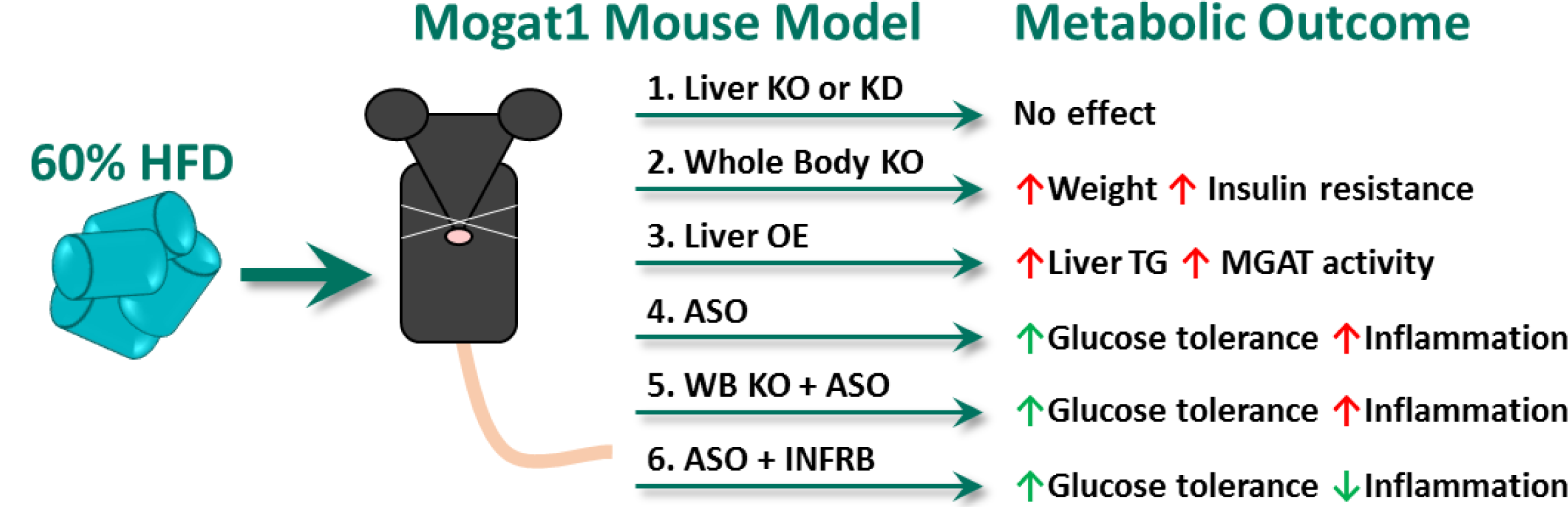

**Highlights:**

- Mogat1 liver-specific KO or KD does not improve metabolism in HFD fed mice.
- Whole-body Mogat1-deletion impairs insulin tolerance in HFD fed mice.
- Mogat1 ASOs improves whole body metabolism independently of gene knockdown.
- Blockade of the INFR response does not prevent off-target effects of Mogat1 ASOs.

## 1. INTRODUCTION

Nonalcoholic fatty liver disease (NAFLD) results from ectopic intrahepatic lipid accumulation (steatosis) and is believed to be a key driver of many metabolic abnormalities associated with obesity including insulin resistance and diabetes [1,2]. In obesity, the liver is overloaded with fatty acids from dietary intake, increased adipose tissue lipolysis, and higher rates of *de novo* lipid synthesis [3–5]. The liver must efficiently store these lipids as neutral triacylglycerol (TAG) to prevent deleterious effects of toxic lipid intermediates such as activation of inflammation, ER stress, and insulin resistance [6].

There are two pathways of TAG synthesis in liver: the glycerol-3-phosphate (G-3-P) pathway and the monoacylglycerol O-acyltransferase (MGAT) pathway. These pathways converge at diacylglycerol (DAG), which is the sole precursor of TAG [7–9]. MGATs acylate monoacylglycerol to form DAG and this enzymatic activity is encoded by several genes including *Mogat1, Mogat2*, and *Dgat1* in mice [10–13]. The importance of MGAT activity in dietary fat absorption by intestinal enterocytes has long been studied [8,14]. In contrast, the G-3-P pathway is considered to be the primary route of TAG synthesis in tissues other than the intestine and the MGAT pathway is believed to be an auxiliary pathway. We have recently demonstrated that *Mogat1* is also highly expressed in adipocytes and may function to suppress aberrant lipolysis [15]. In addition, the expression and activity of the MGATs are increased in humans with NAFLD and mouse models with hepatic steatosis [16–20]. Suppression of hepatic and adipose tissue *Mogat1* expression through use of antisense oligonucleotides (ASO) reduces hepatic MGAT activity and improves hepatic insulin sensitivity and glucose metabolism without reducing hepatic steatosis in obese mice [17,18].

Second generation ASOs are known to target multiple tissues including liver, adipose tissue, and intestine, all of which have MGAT activity [21]. Moreover, off-target effects of ASO treatment have also been described [22]. McCabe and colleagues recently demonstrated an improvement in adipose tissue metabolism by ASO targeting TTC39B mRNA that was independent of target gene knockdown. These effects were mediated through the activation of interferon alpha/beta receptor 1 (IFNAR-1) signaling in adipose-derived macrophages [22]. Thus, we sought to obtain rigorous and independent confirmation that hepatic *Mogat1* plays an important role in obesity-related hepatic insulin resistance and metabolic abnormalities by using novel liver-specific and global *Mogat1* knockout mice. Our data surprisingly suggest that *Mogat1* ASO treatment improves glucose metabolism through *Mogat1*-independent mechanisms and that the effects are not mediated through IFNAR-1 activation.

## 2. METHODS

### 2.1 Generation of mouse models

All mouse studies were approved by the Institutional Animal Care and Use Committee of Washington University. Due to the resistance of female mice to the effects of high fat diet, male mice in the C57BL6/J background were used in all studies. Mice were group housed and maintained on standard laboratory chow on a 12 h light/dark cycle. At eight weeks of age mice were placed on control low-fat diet (LFD, Research Diets, 10 kcal % fat matched sucrose, D12450J) or high-fat diet (HFD, Research Diets, 60 kcal % fat, D12492) for the durations indicated. ASO treatments were performed as previously described [17]. Briefly, starting at 16 weeks on diet, mice were given twice weekly intraperitoneal injections of ASO directed against *Mogat1* (sequence 1: 5’-GATCTTGGCCACGTGGAGAT-3’ (20-mer) or sequence 2: 5’-TGGCCACGTGGAGATACGAT-3” (20-mer) where indicated) or scramble control (Ionis, Pharmaceuticals, Inc., Carlsbad, CA; 25 mg/kg body weight) for three weeks.

Embryonic stem cells used to generate *Mogat1* whole-body knockout and *Mogat1* floxed mice were obtained from the Knockout Mouse Consortium (project# CSD35789) and mice harboring this floxed allele have been previously described [15]. *Mogat1* floxed mice were crossed with mice expressing the Cre recombinase transgene under the control of the albumin promoter (Jackson Laboratory, B6.Cg-Speer6-ps1Tg (Alb-cre) 21Mgn/J). Acute liver-specific knockout mice were generated by retro-orbital injection of *Mogat1* floxed mice with 2.0 ⨯ 10^11^ genomic copies (GC) of adeno-associated virus serotype 8 (AAV8) expressing Cre recombinase under the control of human thyroid hormone-binding globulin (TBG) promoter (Vector Biolabs, VB1724). Control mice received AAV8 expressing enhanced green fluorescent protein (GFP) under control of the same promoter (Vector Biolabs, VB1743). For shRNA studies, C57BL6/J mice (Jackson Laboratory) received retro-orbital injection of either control AAV8-GFP-U6-scramble-shRNA (Vector Biolabs) or AAV8-GFP-U6-*Mogat1*-shRNA (sequences 5-UUUCACCCUCAUGGAAUAUUCGUGCCU-3 and 5-CAAGACGCAAUGUAUGAUUCAAUGGGA-3 [20]; pooled (2.0 x 10^11^ GC total) before injection (Vector Biolabs). For hepatic *Mogat1* overexpression, eight-week-old male C57BL6/J mice were given LFD or HFD for six weeks, then administered AAV8-TBG-eGFP or AAV8-TBG-mouse-*Mogat1* by retro-orbital injection (Vector Biolabs, VB1743, custom refseq# BC106135).

In the IFNAR-1 inhibition study, HFD-fed male C57BL/6J mice were obtained from Jackson Laboratory after 12-weeks of diet. After acclimation and four additional weeks of HFD feeding, mice were weight matched into four treatment groups. Mice were given scramble ASO or ASOs targeting *Mogat1* (5’-TGGCCACGTGGAGATACGAT-3” (20-mer)) as described above. During each ASO treatment mice were also given IP injections of IgG control (BioCell *InVivoI*MAb, #BE0083, clone MOPC-21) or a monoclonal neutralizing antibody targeting interferon alpha/beta receptor 1 (IFNAR-1) (BioCell *InVivoI*MAb, #BE0241, clone MAR1-5A3) [23]. Antibody treatments were given as IP injections of 250 ug per mouse in 100 uL of dilution buffer (BioCell, *InVivo*Pure Ph 6.5 Dilution Buffer, #IP0065) twice a week for the first two weeks of ASO treatment followed by three separate injections of 500 ug per mouse the third week of ASO treatments.

Prior to sacrifice mice were fasted for 4 hours starting at 0900. Mice were euthanized via CO_2_ asphyxiation. Blood was collected from venipuncture of the inferior vena cava into EDTA-coated tubes and plasma was removed by centrifugation. Liver and other tissues were immediately collected, flash-frozen in liquid nitrogen, and stored at -80°C until further use.

### 2.2 Metabolic phenotyping of mouse models

Glucose tolerance tests were performed in mice fasted for 5 hours starting at 0900. Mice were given an intraperitoneal (IP) injection of glucose (1 g/kg body weight dissolved in saline) and blood glucose was measured from the tail using a One Touch Ultra glucometer (Life Scan Inc.) at times indicated. For insulin tolerance tests, mice were fasted for 4 hours before IP injections of recombinant Humulin R® (0.75 U/kg body weight in saline). Tolerance tests were performed one week apart to allow the mice to recover starting at 15 and 16 weeks of diet or week two and three of acute treatments. Body composition was determined in fed mice using ECHO MRI. Because the whole-body null mice had differences in body weights compared to WT mice, these mice were dosed based on lean mass (1 g glucose/kg lean mass and 1.5 U insulin/kg lean mass).

### 2.3 Liver lipids and plasma metabolites

Frozen liver pieces were homogenized with bead disruption in PBS (100 mg ml). Lipids were solubilized in 1 % sodium deoxycholate via vortexing and heating at 37°C for 5 min. Triglycerides were measured enzymatically using the Infinity Triglyceride colorimetric assay kit (Thermo Fisher, TR22421) and normalized to mg tissue (wet weight). Plasma insulin was determined by Singulex Immunoassay (Millipore Sigma) via the Washington University Core Laboratory for Clinical Studies.

### 2.4 mRNA isolation and quantitative PCR

Total liver RNA was isolated from frozen liver samples using RNA STAT (Iso Tech, CS-502) according to the manufacturer’s protocol. RNA was reverse transcribed into cDNA using Taqman high capacity reverse transcriptase (Life Technologies, 43038228). Quantitative PCR was performed using Power SYBR green (Applied Biosystems, 4367659) and measured on an ABI PRISM 7500 or ABI QuantStudio 3 sequence detection system (Applied Biosystems). Results were quantified using the 2^-ΔΔCt^ method and shown as arbitrary units relative to control groups. Primer sequences are listed in supplemental table 1.

### 2.5 Glycogen concentration analyses

Samples of frozen tissues (30 to 90 mg) were hydrolyzed in 0.3 ml of 30% (wt/vol) KOH solution in a boiling water bath for 30 min. At 10 and 20 min of the incubation, tubes were vortexed to facilitate digestion. After cooling to room temperature, 0.1 ml of 1 M Na2SO4 and 0.8 ml of ethanol were added, the samples were then boiled again for 5 min to facilitate precipitation of glycogen and then centrifuged at 10,000 x g for 5 min. The glycogen pellet was dissolved in 0.2 ml of water, and two additional ethanol precipitations were performed. The final pellet was dried and dissolved in 0.2 ml of 0.3 mg/ml amyloglucosidase (Sigma Aldrich 11202332001) in 0.2 M sodium acetate buffer (pH 4.8) and incubated for 3 h at 40°C. The reaction mixture was diluted two- to fivefold with water. To determine the glucose concentration, 5 ml of the diluted sample was added to 0.2 ml of the glucose assay solution (0.3 M triethanolamine–KOH (pH 7.5), 1 mM ATP (Sigma Aldrich, A6559), 0.9 mM b-NADP (Sigma Aldrich, 10128031001), and 5 mg/ml of G-6P dehydrogenase (Sigma Aldrich, 10165875001)). The absorbance at 340 nm was determined before and after addition of 1 mg of hexokinase (Sigma Aldrich, 11426362001). Glycogen content is expressed as micromoles of glucosyl units per gram (wet weight).

### 2.6 Primary hepatocyte isolations

Primary hepatocytes were isolated as previously described [24]. Briefly, 10-16-week-old female mice were given an overdose of isoflurane prior to perfusions. The livers were perfused via catheterization of the hepatic portal vein and flushed with 30 ml of HBSS (Ca^2+^/Mg^2+^-free, 0.5 mM EGTA) prior to digestion with 20 ml of collagenase solution (Type IV collagenase (sigma S5138) at 1 mg/ml in DMEM (serum free, 1 mM sodium pyruvate). Following perfusion, the livers were removed and disrupted in the collagenase solution. Cells were added to ice-cold complete DMEM (10% FBS, 1 mM sodium pyruvate, 100 U PenStrep, 0.25 µg/ml amphotericin b) and passed through a 50 µM filter prior to centrifugation at 50 g x 2 min. The supernatant was removed and hepatocytes were resuspended in complete DMEM and washed two more times. 500,000 cells were then directly added to RNA STAT for RNA isolation and quantification.

### 2.7 MGAT enzymatic assay

MGAT activity was determined as previously described [17]. Liver (50 mg/ml) was homogenized by probe sonication in ice-cold membrane buffer (50 mM Tris-HCl pH 7.4, 1 mM EDTA, 250 mM sucrose, and complete protease inhibitors table (Roche Diagnostics, A32965)). Homogenates were clear of whole-cell debris by low speed centrifugation at 500 g x 10 min at 4°C. The resulting supernatants were spun at 100,000 g x 60 min at 4°C using a Beckman benchtop ultra-centrifuge. The cytosolic fractions (supernatant) were removed and the membranes were reconstituted via pipetting in membrane buffer without protease inhibitors. Proteins were quantified using the bicinchoninic acid (BCA) assay according to the manufacture’s protocols (Themo Fisher, 23225). Fifty micrograms of membrane were incubated in assay buffer (5 mM MgCl2, 1.25 mg/ml BSA, 200 mM sucrose, 100 mM Tris-HCl (pH 7.4)) containing 20 µM [^14^C]oleoyl-CoA (American Radiolabeled Chemicals, ARC 0527), and 200 µM sn-2-oleoylglycerol (Caymen Chemical, 16537) for 10 min. The reaction was stopped with 50 µl 1% phosphoric acid. Lipids were extracted in 2:1 v/v % CHCl3:MeOH and separated by thin-layer chromatography in hexane/ethyl ether/acetic acid (80:20:1, v /v/v %). Samples were run against standards for oleic acid, 1,3 diacylglycerol and triglyceride and corresponding spots were scraped from the plate and ^14^C-radioactivity was measured via scintillation counter. Backgrounds were calculated from reaction mixtures without membrane fractions.

### 2.8 Western blotting

Membrane and cytosolic protein samples were isolated as described in (Section 2.7). Fifty micrograms of protein were loaded onto 4-15% acrylamide pre-cast gels (BioRad, 64329760) in Lamellae loading buffer (BioRad, 161-0737). Proteins were transferred to PVDF membranes in Tris-glycine buffer with 10% methanol. Following transfer, membranes were blocked in 5% BSA in TBS for 1 hour prior to overnight antibody incubations in 5% BSA in TBS. Antibodies used are as follows: *Mogat1* (Santa Cruz Biotechnologies, 32387, 1:500) green fluorescent protein (GFP, Cell Signaling Technologies, 2555, 1:1000), Calnexin (Enzo Life Sciences, ADI-SPA-860, 1:1000) and mouse γ-tubulin (Sigma-Aldrich, T6557, 1:1000). Following incubations, membranes were washed and incubated in secondary antibodies (LiCor) prior to imaging on a LiCor Odyssey.

### 2.9 Statistical analysis

Data were analyzed using GraphPad Prism software. Independent and paired T-tests, one-way analysis of variance (ANOVA), or factorial ANOVAs were performed where appropriate. Secondary post-hoc analysis found differences in groups using either Tukey or Sidek multiple comparisons were appropriate. *P* < 0.05 was considered significant.

## 3. RESULTS

### 3.1 Liver-specific knockout of *Mogat1*

We generated mice with hepatocyte-specific deletion of *Mogat1* by crossing *Mogat1* floxed mice [15] to homozygosity with mice expressing albumin promoter-driven Cre recombinase (Figure 1). The resulting offspring mice were outwardly normal on a standard chow diet. When given ad libitum access to a 10% fat (LFD) or 60% fat (HFD), the mice gained weight similar to littermate fl/fl controls (Figure 1A). Contrary our previous studies showing a beneficial effect of *Mogat1* ASO on metabolic parameters in obese mice, glucose and insulin tolerance was not different between genotypes on LFD or HFD (Figure 1B, C). We confirmed that *Mogat1* expression was markedly suppressed in *Mogat1* fl/fl Alb-Cre+ livers and freshly isolated primary hepatocytes (of note, *Mogat1* expression drastically decreases after hepatocyte isolation and plating, data not shown) (Figure 1D, E). *Mogat2* tended to be, but was not significantly increased in knockout mice or isolated hepatocytes (Figure 1F, G). Liver weight was not altered by *Mogat1* knockout (Figure 1H). Hepatic TAG content increased with HFD but was similar between genotypes (Figure 1I). In contrast to previous work with *Mogat1* ASO-treated mice, MGAT activity in isolated liver membranes was not affected by *Mogat1* knockout (Figure 1J and ref. [17]).

**Figure 1.**
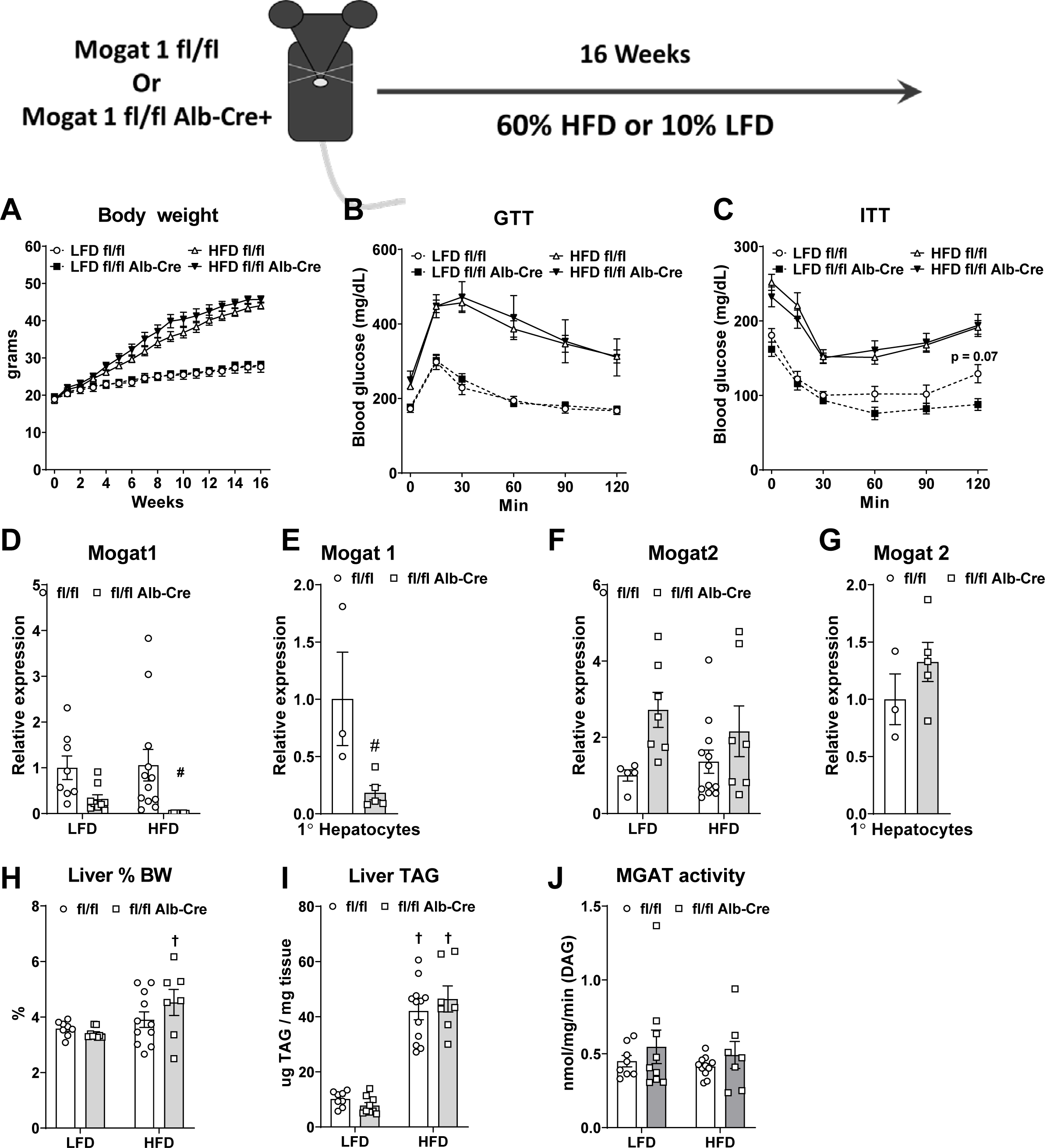
Constitutive liver-specific Mogat1 deletion does not improve insulin sensitivity in mice. Male Mogat1 fl/fl mice and littermate Mogat1 fl/fl albumin Cre+ mice were fed a LFD or a HFD starting at eight weeks of age for 16 weeks. Mice were fasted for 4 hours prior to sacrifice and tissue collection. A: HFD increased bodyweight in both groups. B,C: HFD fed mice have impaired glucose and insulin tolerance compare to LFD groups. D-G: Mogat1 gene expression is reduced in knockout liver and primary hepatocytes without significant compensation of Mogat2. H,I: HFD increased liver weight and TAG content in both genotypes. J: MGAT activity was not affected by diet or genotype. Data are expressed as means ± S.E.M. # *p* < 0.05 gene effect, † *p* < 0.05 diet effect; *n* = 5-10 for mouse studies, *n* = 3-5 female mice for primary hepatocyte isolations.

### 3.2 AAV8-mediated liver-specific knockout of *Mogat1*

To avoid possible compensation from chronic *Mogat1* knockout, we injected *Mogat1* fl/fl mice with an AAV8-TBG-Cre or eGFP control after 16 weeks of HFD feeding (Figure 2). After three weeks, AAV8-Cre knockout mice had similar body weight and no differences in glucose or insulin tolerance (Figure 2A-C). The AAV8-Cre reduced Mogat 1 expression but did not affect *Mogat2* expression (Figure 2D). Similar to constitutive *Mogat1* liver-specific knockout, acute deletion of hepatic *Mogat1* did not alter liver weight, TAG content, and actually, slightly increased MGAT activity (Figure 2E-G). Together, these data indicate that genetic deletion of *Mogat1* in liver of diet-induced obese mice does not affect glucose or insulin tolerance, and suggests compensation from other enzymes with MGAT activity [11,12,25].

**Figure 2.**
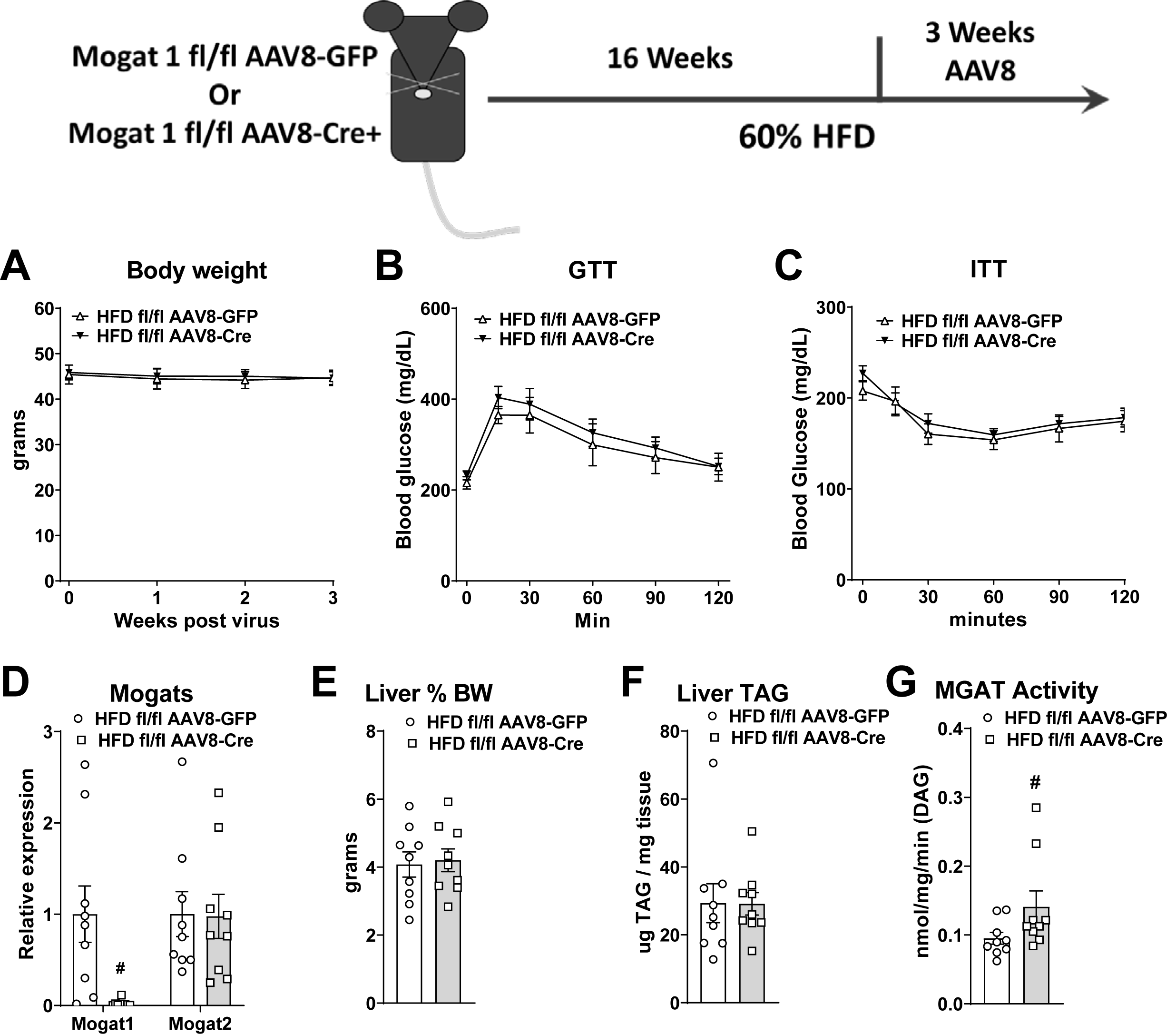
Acute liver-specific deletion of Mogat1 does not improve glucose or insulin tolerance in HFD fed mice. Male Mogat1 fl/fl mice were fed a HFD starting at eight weeks of age. After 16 weeks mice were given a retro-orbital injection of AAV8-TBG-eGFP or Cre recombinase (2 x 10^11^ GC per mouse) and remained on diet for an additional 3 weeks. Mice were fasted for 4 hours prior to sacrifice and tissue collection. A: Mogat1 knockdown did not affect body weight. B,C: Acute liver-specific knockout did not improve glucose or insulin tolerance. D: Mogat1 knockout reduced Mogat1 expression without increasing Mogat2 expression in liver. E-F: Liver weights and TAG content were not difference among groups. G: Hepatic MGAT activity was not reduced by acute Mogat1 knockdown in liver. Data are expressed as means ± S.E.M. # *p* < 0.05 gene effect; *n* = 9.

### 3.3 *Mogat1* whole-body deletion

*Mogat1* ASO treatment likely results in gene knockdown in multiple tissues [21]. Thus, we generated *Mogat1* whole body null mice (MOKO) (Figure 3). MOKO mice were outwardly normal on standard chow diet. Unexpectedly, when given a HFD, MOKO mice gained more weight than littermate wild-type control mice (Figure 3A, B). The heterozygous littermates had an intermediate phenotype (Figure 3B and data not shown). The increase in weight gain was not attributed to increased adiposity as evident by fat mass and percent adiposity in ECHO MRI and individual fat pad weights and thus may be due to larger body size (Figure 3B). Glucose tolerance was not affected by *Mogat1* knockout; however, HFD-fed MOKO mice had reduced insulin tolerance and increased plasma insulin concentrations, which could be indicative of insulin resistance (Figure 3D, E). *Mogat1* gene expression was undetectable in livers of MOKO mice (Figure 3F) or in any other tissue analyzed (data not shown). Lastly, MOKO mice had larger livers compared to wild-type littermates, but similar TAG content and MGAT activity (Figure 3G-I).

**Figure 3.**
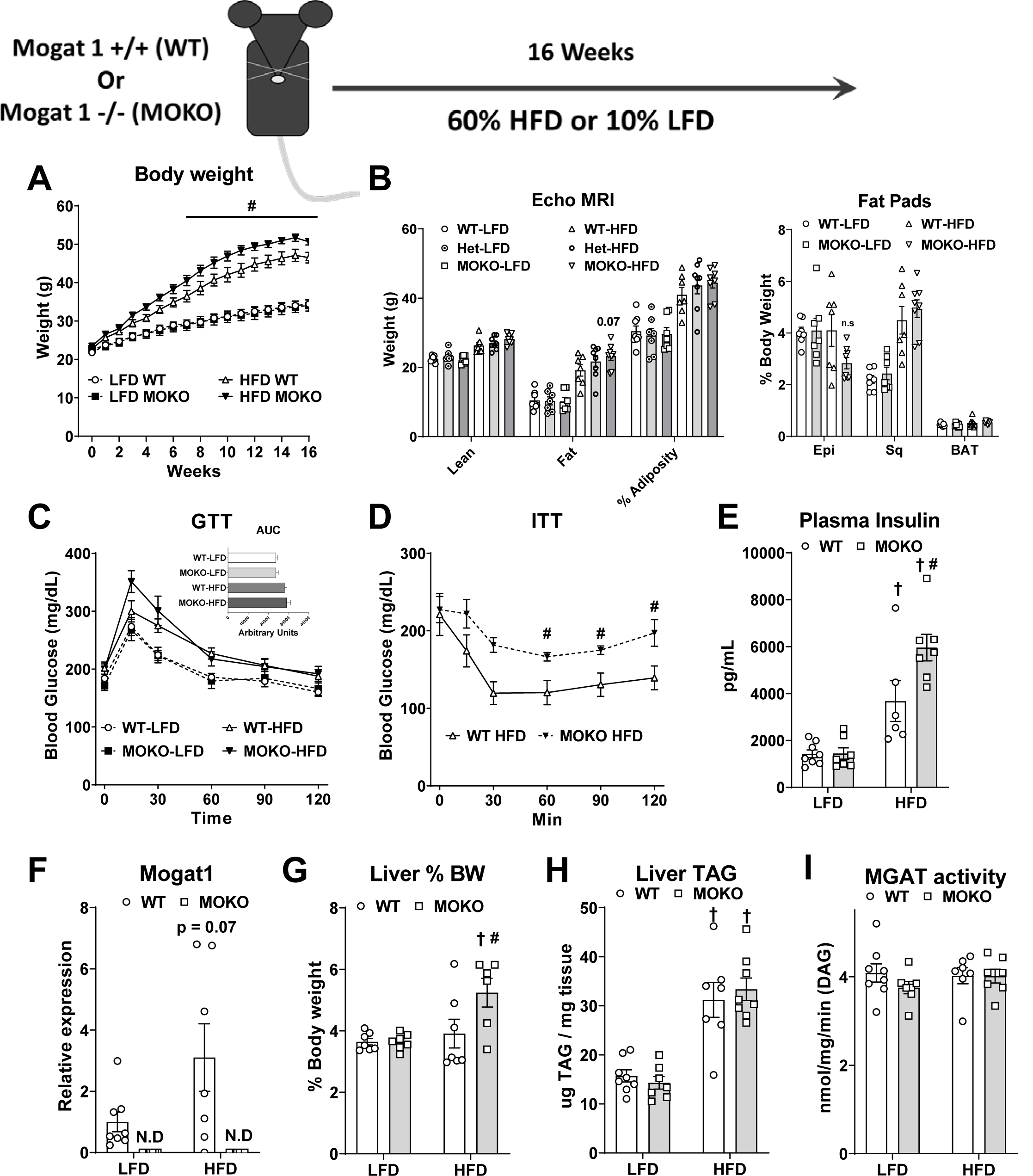
Whole-body deletion of Mogat1 causes weight gain and insulin intolerance on a HFD. Male wild-type (WT) and littermate Mogat1 whole body knockout (MOKO) mice were fed a LFD or a HFD starting at eight weeks of age for 16 weeks. Mice were fasted for 4 hours prior to sacrifice and tissue collection. A: Mogat1 knockout mice gain more weight on a HFD than littermate WT controls. B: ECHO MRI indicates MOKO mice have increased whole body mass, while the heterozygous mice had an intermediate phenotype compared to WT controls. C: Glucose tolerance (dosed on lean mass) was not significantly change in MOKO mice. D,E: HFD fed MOKO mice had significantly impaired insulin tolerance and plasma insulin levels compared to WT controls. F: Mogat1 gene expression was not detectable in MOKO mice. G,H: Liver weight but not TAG was increased in MOKO mice fed a HFD compared to LFD controls. I: MGAT activity was unaffected by either diet or genotype. Data are expressed as means ± S.E.M. # *p* < 0.05 gene effect, † *p* < 0.05 diet effect; *n* = 6-7.

### 3.4 Hepatic *Mogat1* overexpression

We next wanted to determine if overexpression of hepatic *Mogat1* in the context of LFD or HFD was sufficient to promote hepatic steatosis, insulin resistance, as well as deleterious effects on systemic glucose metabolism. We treated C57BL6/J mice with LFD or HFD for 6 weeks then injected them with AAV8 expressing either eGFP or mouse *Mogat1* under control of the hepatocyte-specific TBG promoter (Figure 4). *Mogat1* overexpressing (OE) mice gained similar weight as the GFP control-treated mice on either diet (Figure 4A). After 16 weeks of diet (10 weeks of overexpression), there was no difference in glucose or insulin tolerance between groups on HFD (Figure 4B, C). *Mogat1* and GFP expression was significantly increased by the administration of each AAV8, respectively and there was no difference in *Mogat2* expression between groups (Figure 4D-F). *Mogat1* overexpression did not alter liver weight but did increase liver TAG and MGAT activity in the LFD groups (Figure 4H, I). *Mogat1* overexpression did not affect MGAT activity in HFD fed mice, likely due to the effect of HFD in increasing MGAT activity. *Mogat1* protein expression was restricted to liver membranes where MGAT enzymatic activity occurs (Figure 4J). Thus, we conclude that hepatic *Mogat1* overexpression is sufficient to induce hepatic steatosis on a LFD, but not insulin resistance or systemic dysregulation of glucose metabolism.

**Figure 4.**
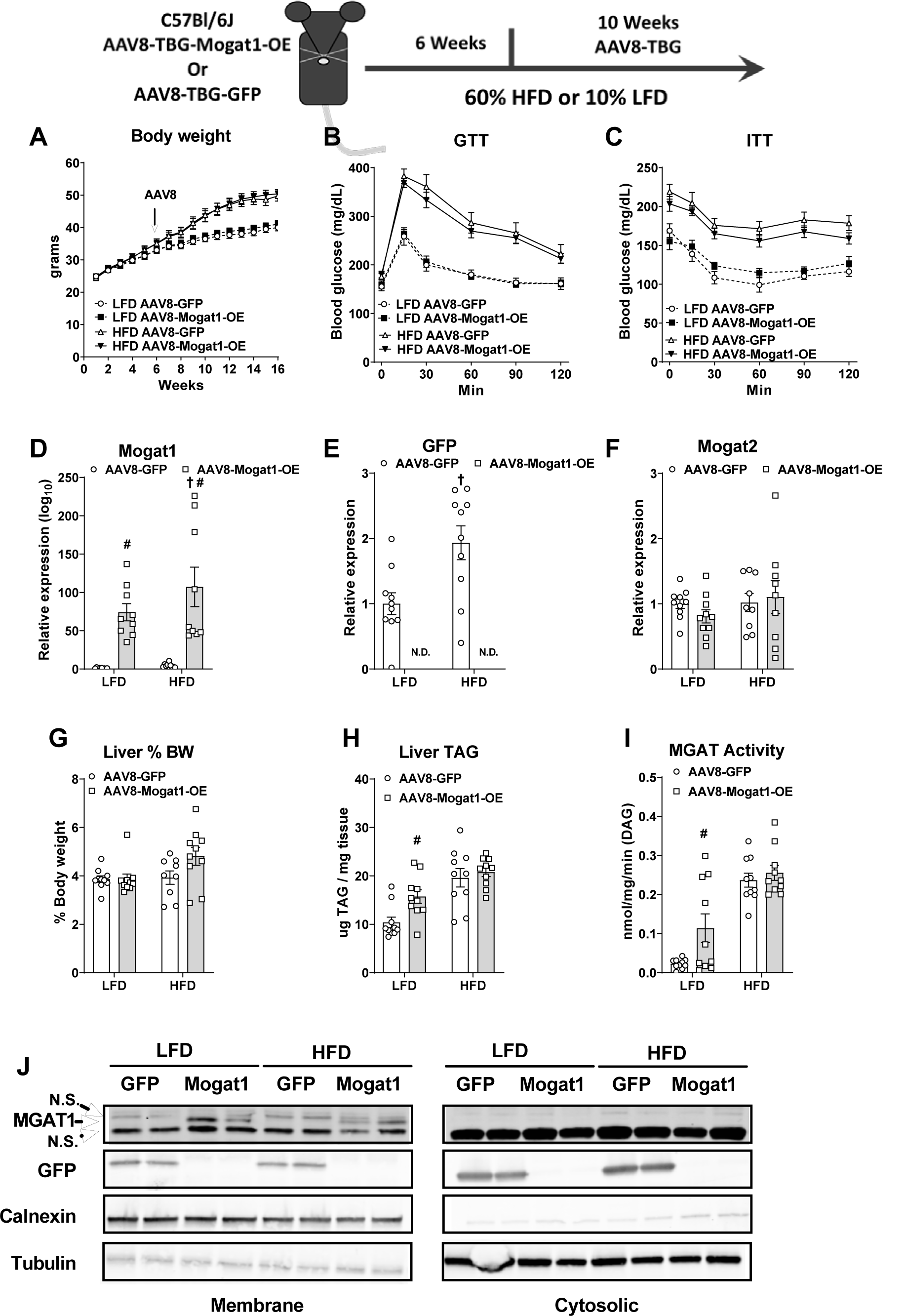
Hepatic Mogat1 overexpression increases liver TAG and MGAT activity in mice fed a LFD. Male C57BL6/J mice were fed LFD or a HFD starting at eight weeks of age. After 6 weeks of diet mice were injected (retro-orbitally) with AAV8-TGB-eGFP- or AAV8-TGB-Mogat1 (2 x 10^11^ GC per mouse) and remained on diet for an additional 10 weeks. Mice were fasted for 4 hours prior to sacrifice and tissue collection. A: HFD fed mice gained weight compare to LFD fed mice in both treatment groups. B,C: Mogat1 overexpression did not impair glucose or insulin tolerance. D-F: Mogat1 and GFP gene expression was significantly increased in AAV8-Mogat1 and AAV8-GFP treated mice, respectively without a change in Mogat2 gene expression. G: Liver weight was unaffected by any treatment. H,I: AAV8-Mogat1 overexpression increased both liver TAG and MGAT activity in LFD mice. J: Western blot analysis indicated Mogat1 and eGFP protein increased in membrane fractions of AAV8 treated livers. Data are expressed as means ± S.E.M. # *p* < 0.05 gene effect, † *p* < 0.05 diet effect; *n* = 8-10.

### 3.5 shRNA mediated knockdown of *Mogat1*

Work conducted independently of our lab has shown that adenovirus-delivered *Mogat1* shRNA to knockdown hepatic *Mogat1* expression decreased hepatic triglycerides and plasma glucose in mouse models of fatty-liver disease [26,20]. As an alternative to our genetic models, we used AAV8 to deliver the same shRNAs targeted against *Mogat1* or scramble controls in HFD-fed C57BL6/J mice (Figure 5). However, *Mogat1* knockdown by shRNA did not improve glucose or insulin tolerance in HFD fed mice in our study (Figure 5A-C). *Mogat1* gene expression was significantly reduced by *Mogat1* shRNA treatment, but was not returned to levels of LFD mice, and *Mogat1* shRNA did not affect liver weight or TAG levels (Figure 5D-G). These data suggest that shRNA-mediated *Mogat1* knockdown does not improve insulin sensitivity or glucose metabolism.

**Figure 5.**
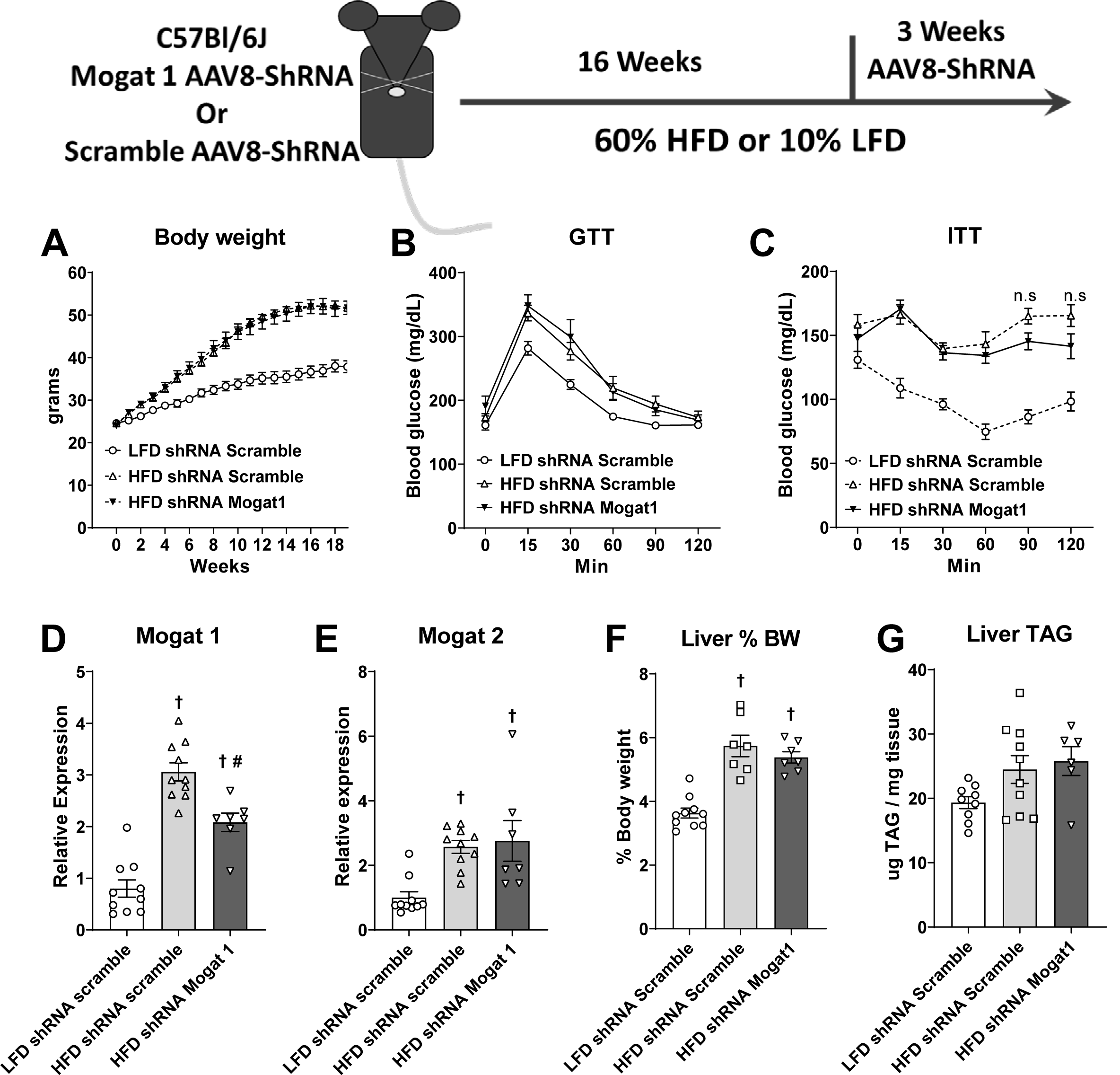
shRNA mediated Mogat1 knockdown did not improve glucose or insulin tolerance on a HFD. Male C57Bl/6J mice were fed a 10% LFD or a 60% HFD starting at eight weeks of age. After 16 weeks of diet mice were injected (retro-orbitally) with AAV8-U6-shRNA Scramble or shRNAs targeted against Mogat1 (2 x 10^11^ GC per mouse) and remained on diet for an additional three weeks. Mice were fasted for 4 hours prior to sacrifice and tissue collection. A: HFD fed mice gained weight compare to LFD Scramble treated controls. B,C: HFD fed mice had impaired glucose and insulin tolerance compared to LFD Scramble control treated mice. D,E: Liver weight and TAG were increased by the HFD but unaffected by Mogat1 shRNA treatment. F,G: Mogat1 ShRNA treatment reduced Mogat1 expression but did not affect Mogat2 expression. Data are expressed as means ± S.E.M. # *p* < 0.05 from HFD shRNA Scramble control mice, † *P* < 0.05 from LFD shRNA Scramble controls; *n* = 7-10.

### 3.6 *Mogat1* ASO treatment improves hepatic insulin sensitivity on HFD

To confirm our previous reports indicating that *Mogat1* suppression by ASO improves hepatic insulin sensitivity [17,18], we fed C57BL6/J mice with a LFD or HFD. After 16 weeks, mice were weight-matched again and randomized to receive ASOs targeting either *Mogat1* or scramble control (Figure 6). As previously described [17], *Mogat1* ASO treatment lowered blood glucose and plasma insulin concentrations compared to scramble treated controls in HFD fed mice without affecting body weight (Figure 6A-C). Hepatic *Mogat1* and *Mogat2* gene expression were significantly increased by HFD, but *Mogat1* ASO treatment suppressed *Mogat1* expression without affecting *Mogat2* (Figure 6D). *Mogat1* ASO treatment increased liver weight (Figure 6E) without increasing TAG content (Figure 6F). The increased liver weight could be due to glycogen accumulation, which was higher in *Mogat1* ASO treated mice (Figure 6G). Together these data confirm our previous reports that *Mogat1* ASO treatment increases improves hepatic metabolism.

**Figure 6.**
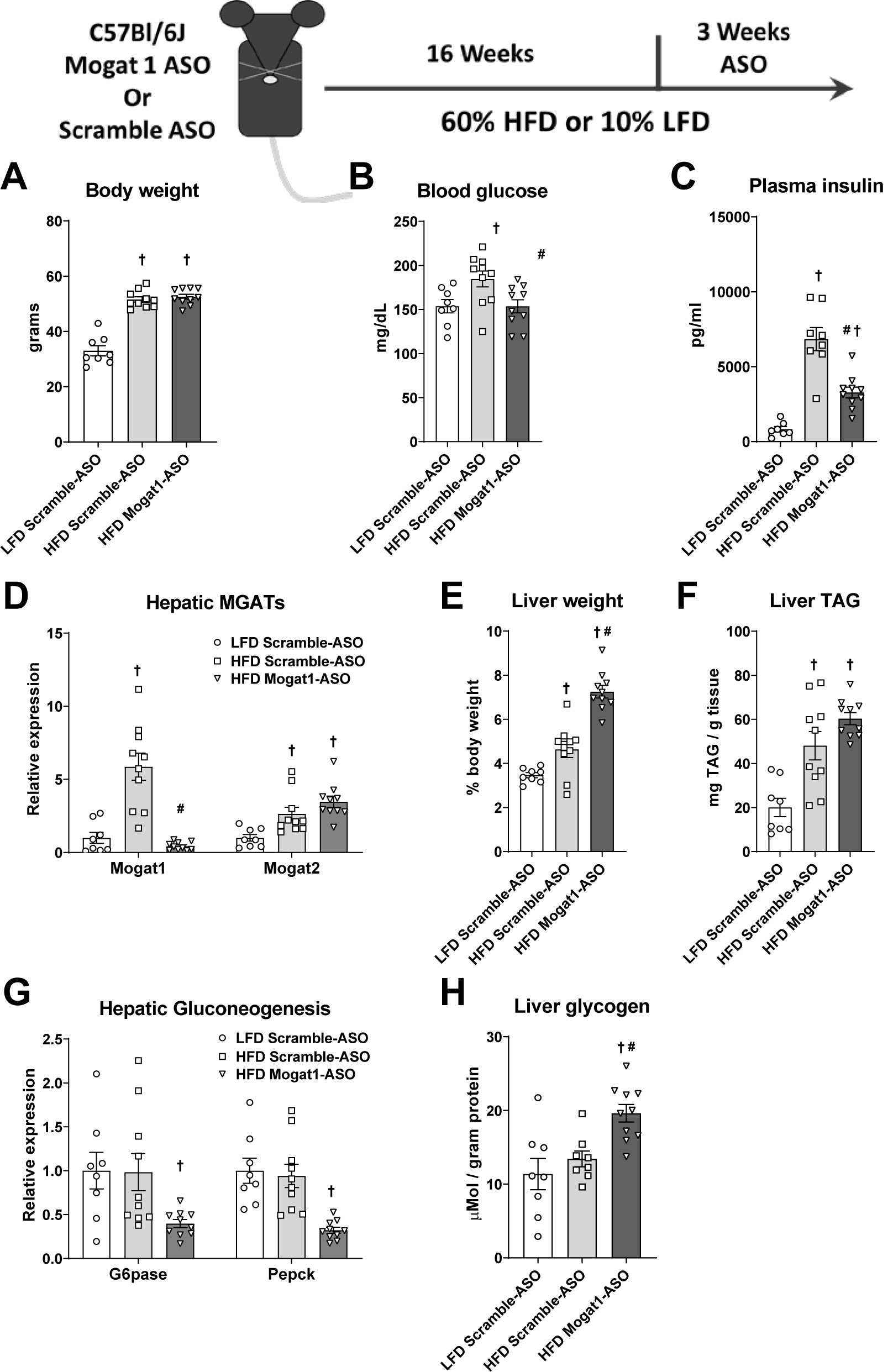
Mogat1 antisense oligonucleotide (ASO, sequence 1) treatment improves insulin sensitivity in HFD fed mice. Male C57Bl/6J mice were fed a LFD or a HFD starting at eight weeks of age. After 16 weeks of diet, mice were injected (intraperitoneally) twice weekly with ASOs targeted against Mogat1 or scramble control (25 mg/Kg) for three weeks. Mice were fasted for 4 hours prior to sacrifice and tissue collection. A: HFD fed mice gained more weight compare to LFD fed controls. B,C: Control ASO treated mice had significantly higher blood glucose and plasma insulin than LFD Control ASO treated mice and HFD Mogat1 ASO treated mice. D: Mogat1 ASO treatment significantly reduced hepatic expression of Mogat1, but not Mogat2 as measured by RT-qPCR. E: Mogat1 ASO treatment increased liver weight (%BW) on HFD. F: Liver triglycerides (TAG) were measured enzymatically and were increased by the HFD. G: Hepatic gene expression of Gpase, Pepck, and Pygl were reduced in the HFD Mogat1 ASO treated mice. H: Glycogen was extracted and measure enzymatically and increased in HFD Mogat1 ASO treated mice. Data are expressed as means ± S.E.M. # *p* < 0.05 ASO effect, † *p* < 0.05 diet effect; *n* = 7-10.

### 3.7 *Mogat1* ASO treatment improves glucose tolerance in *Mogat1* whole-body null mice

Given the inability to reproduce the insulin-sensitizing effects of *Mogat1* ASOs in any of our genetic or shRNA *Mogat1* deletion models, we tested whether *Mogat1* ASO treatment improves glucose metabolism independently of *Mogat1*. We treated MOKO mice, fed a HFD for 16 weeks, with ASOs targeting *Mogat1* or scramble controls (Figure 7). Remarkably, *Mogat1* ASO treatment improved glucose tolerance in both MOKO and WT mice (Figure 7A, B). Furthermore, *Mogat1* ASO treatment increased liver weight and liver TAG in WT mice, with a trend in the MOKO groups (Figure 7C, D). *Mogat1* ASO treatment suppressed *Mogat1* expression in WT mice (Figure 7E).

**Figure 7.**
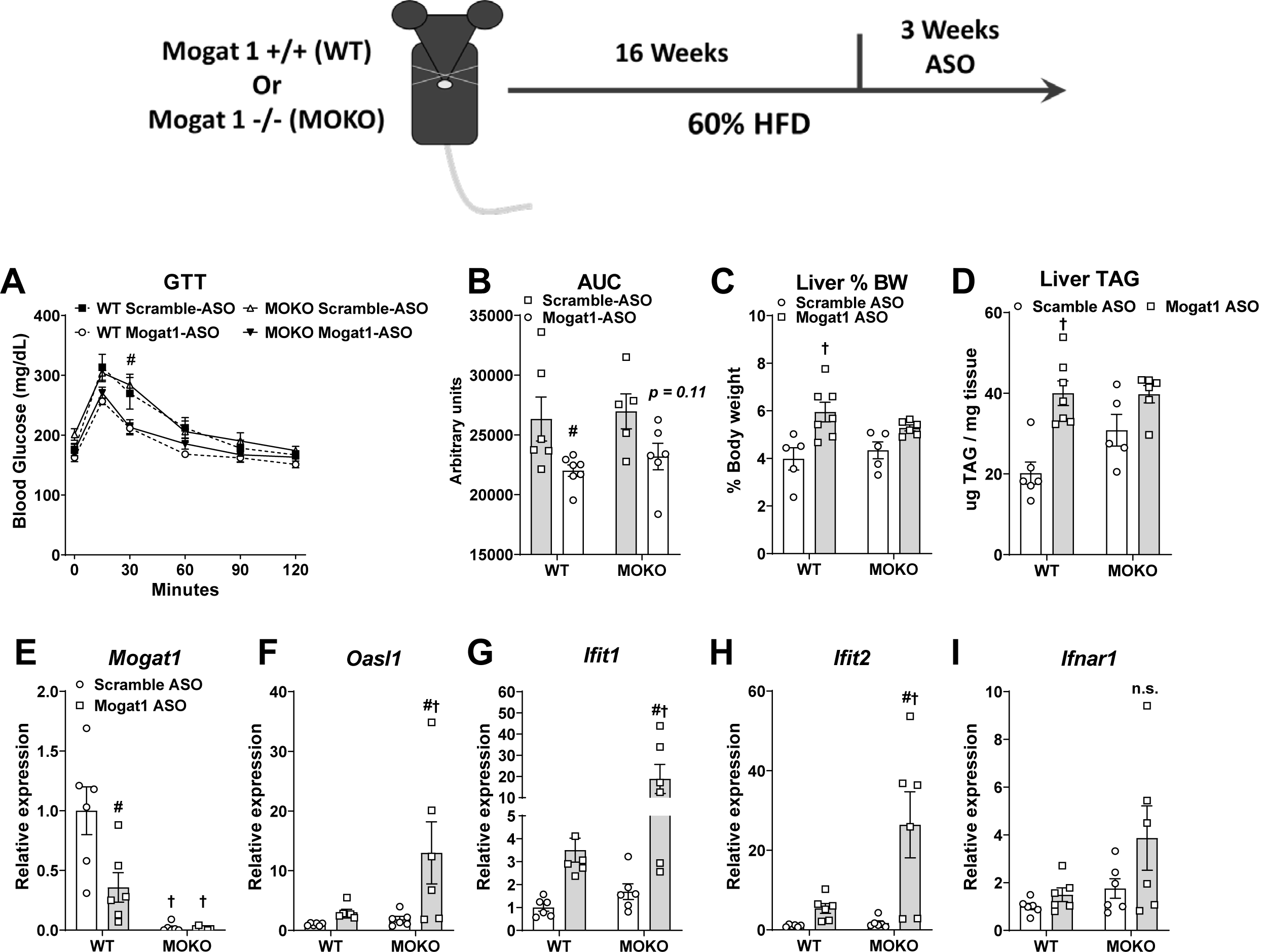
Mogat1 ASO treatment improves glucose tolerance in whole-body Mogat1 null mice on a HFD. Male wild-type (WT) and littermate Mogat1 whole body knockout (MOKO) mice were fed a HFD starting at eight weeks of age. After 16 weeks of diet, mice were injected (intraperitoneally) twice weekly with ASOs targeted against Mogat1 or scramble control (25 mg/Kg) for three weeks. Mice were fasted for 4 hours prior to sacrifice and tissue collection. A,B: Mogat1 ASO treatment improves glucose tolerance despite Mogat1 expression. C,D: Mogat1 ASO treatment causes hepatomegaly and increases liver TAG in wild-type mice on HFD. E: Mogat1 gene expression was reduced by ASO treatment and nearly absent in MOKO livers. F-I: Gene expression markers of interferon signaling are increased in Mogat1 ASO treated MOKO mice on HFD. Data are expressed as means ± S.E.M. # *p* < 0.05 from Scramble ASO † *p* < 0.05 from WT controls; *n* = 5-7.

Recently, McCabe et al. demonstrated that ASO targeting TTC39B non-specifically protects against diet-induced obesity and induces adipose tissue browning through activation of a type I interferon receptor (IFNAR-1) in adipose tissue-derived macrophages, even in TTC39B knockout mice [22]. Although we did not observe any effect on body weight associated with use of the *Mogat1* ASO, we found that expression of several key indicators of IFNAR-1 signaling (*Oasl1, Ifit1*, and *IIfit2*) were significantly increased in liver of MOKO mice treated with Mogat 1 ASOs (Figure 7F-H). The *Mogat1* ASO also tended to increase the expression of these genes in WT mice, while *Ifnar1* gene expression was unchanged in all mice (Figure 7I).

### 3.8 IFNAR-1 blockade does not prevent improvements in glucose homeostasis in *Mogat1* ASO treated mice

To determine if the increase in IFNAR-1 activation in response to *Mogat1* ASO treatment drives the improvements in glucose metabolism, we injected ASO-treated mice with an IFNAR-1 neutralizing antibody or IgG control during a HFD challenge (Figure 8)[22,27]. We repeated the previous observation that the *Mogat1* ASO improved glucose tolerance in HFD fed mice (Figure 8A, B), but co-treatment with IFNAR-1 neutralizing antibody did not block the improvements in glucose tolerance in *Mogat1* ASO treated mice (Figure 8A, B). Liver weight and TAG content were unaffected by any treatment (Figure 8C, D). *Mogat1* ASO treatment suppressed liver *Mogat1* gene expression in both IgG and IFNAR-1 neutralizing antibody treated mice (Figure 8E). *Mogat1* ASO again activated IFNAR-1 responsive genes in liver, but each of these effects was blocked by use of the IFNAR-1 neutralizing antibody (Figure 8F-I). *Mogat1* ASO treatment also suppressed *Mogat1* gene expression (Figure 8J) and increased *Ifitl2* gene expression but not *Ifitl1, Oasl1*, and *Ifnar1* in adipose tissue (Figure 8K-N). Lastly, gene expression of the WAT beiging marker, *Ucp1* was induced by *Mogat1* ASO and blocked by IFNAR-1-Ab (Figure 8O). Other markers of WAT beiging were unaffected by either treatment (*Ppargc1a* or *Arb2*) (Figure 8P-Q). These data indicate that *Mogat1* ASO treatments non-specifically improve glucose metabolism, but this effect is not mediated through the previously implicated mechanism of IFNAR-1 activation [22].

**Figure 8.**
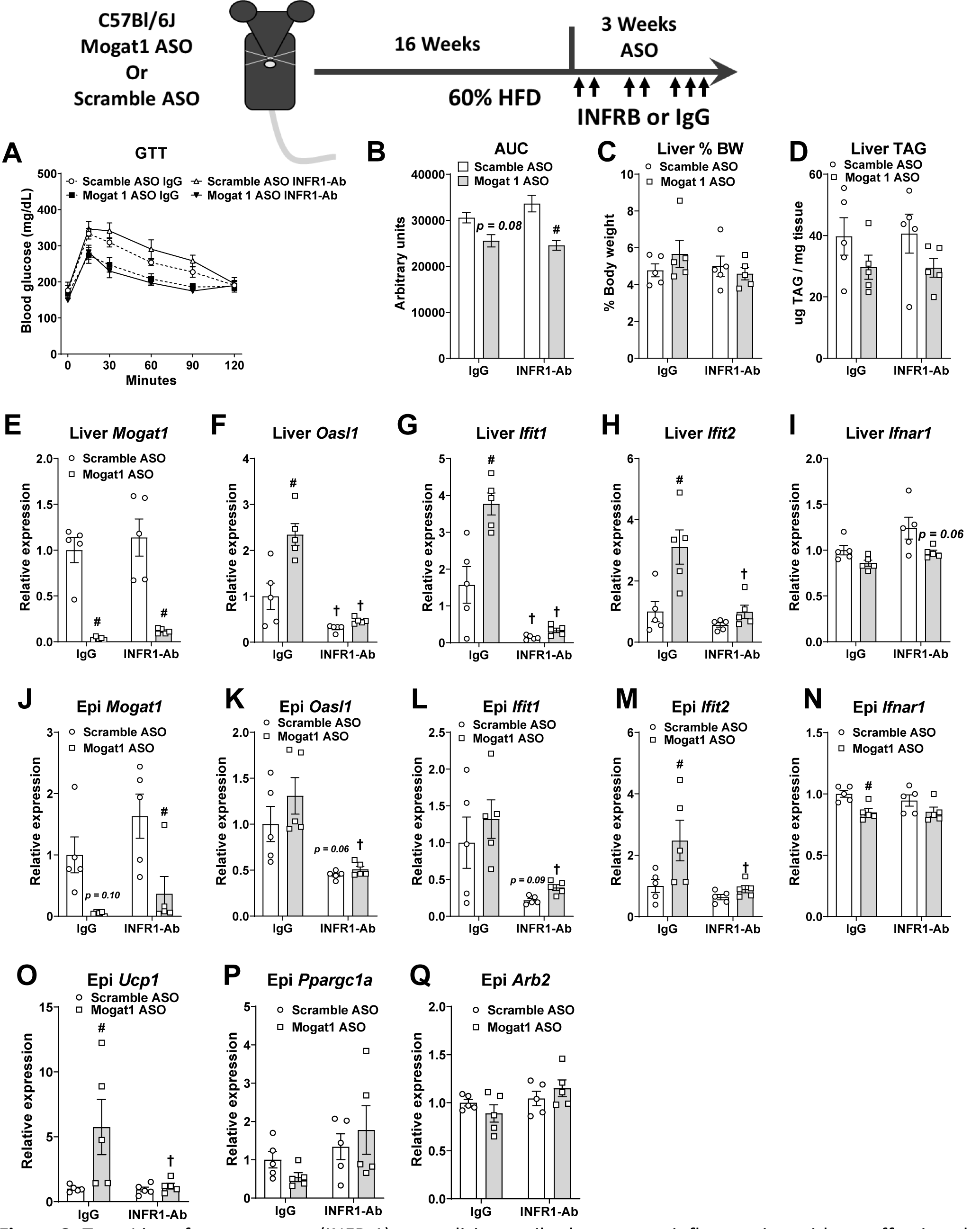
Type I interferon receptor (INFR-1) neutralizing antibody prevents inflammation without effecting glucose tolerance in mice fed a HFD. Male C57BL/6J mice were fed a HFD starting at eight weeks of age. After 16 weeks of diet, mice were injected (intraperitoneally) twice weekly with ASOs targeted against Mogat1 (sequence 2) or scramble control (25 mg/Kg) with an INFR-1 neutralizing antibody (INFR-Ab) or IgG control (250 ug per mouse twice a week for weeks 1&2, and 500 ug per mouse 3 times for week 3). Mice were fasted for 4 hours prior to sacrifice and tissue collection. A,B: Mogat1 ASO treatment improves glucose tolerance despite INFR-1 antibody treatment. C,D: Liver weight and TAG were unaffected by any treatment E: Liver Mogat1 gene expression was reduced by Mogat1 ASO treatment compared to controls. F-I: Liver gene expression markers of interferon signaling are increased in Mogat1 ASO treated mice and inhibited by INFR-Ab. J: Epididymal Mogat1 gene expression was decreased by Mogat1 ASO treatment. K-N: Epididymal gene expression markers of interferon signaling are inhibited by INFR-Ab. O-Q: Expression of adipose beiging genes were increased by Mogat1 ASO and inhibited by INFR-Ab treatment. Data are expressed as means ± S.E.M. # *p* < 0.05 from Scramble ASO † *p* < 0.05 from IgG controls; *n* = 5.

## 4. DISCUSSION

Hepatic steatosis is tightly linked to insulin resistance and development of systemic metabolic abnormalities. Previous work by our group and others using RNA interference approaches has demonstrated that targeting *Mogat1* in livers of obese mice improves glucose homeostasis and insulin resistance [17,18,20,26]. In the present study, we knocked out *Mogat1* specifically in hepatocytes, both chronically and acutely, and also generated a global *Mogat1* knockout, but did not observe any improvements in glucose homeostasis or insulin sensitivity in diet-induced obese mice. Hepatic *Mogat1* overexpression was sufficient to increase hepatic MGAT activity and TAG content, but only in LFD-fed mice, and did not result in insulin resistance or glucose intolerance. Although we reproduced our previous findings to demonstrate that *Mogat1* ASO improves glucose tolerance in obese mice, we also conclusively demonstrate that this ASO still improves metabolic homeostasis in mice with global *Mogat1* deletion. Lastly, we confirm these improvements were not due to activation of interferon signaling through IFNAR-1 as recently reported for an ASO against TTC39B [22]. Collectively, these data suggest that *Mogat1* ASO treatment improves glucose homeostasis and insulin sensitivity independently of knocking down *Mogat1* expression and by mechanisms that remain to be elucidated.

The development of technologies for in vivo RNA interference through use of modified oligonucleotides has been a boon to hepatology and related disciplines. The ease of administration and liver-trophic nature of these designer oligonucleotides has allowed researchers to suppress the expression of a variety of genes in the liver and also led to clinically-approved approaches to treating metabolic disease [28,29]. We were quite flummoxed when characterizing the phenotype of mice lacking *Mogat1* in liver by the inability to phenocopy our previously reported phenotypes obtained by using ASOs [17,18,30] or adenoviral-driven expression of shRNA [20,26]. We have considered a number of possibilities to explain this. It is likely that genetic deletion of *Mogat1* leads to compensatory changes in other enzymes that have MGAT activity, since ASO treatment leads to reduced MGAT activity in liver [17] whereas MGAT knockout either acutely or chronically does not (Figures 1-3). Another possibility is that the *Mogat1* ASO was mediating its effects on a tissue other than liver. Brandon and colleagues recently reported that that liver-specific KO of protein kinase C ε (PKCε) did not phenocopy previous ASO work conducted in rats and found instead that deletion of PKCε in adipose tissue, which is often targeted by ASOs as well, produced an insulin sensitive phenotype [31,32]. However, our studies with adipocyte-specific *Mogat1* KO mice (manuscript in preparation) or with the global MOKO mice did not reveal an insulin sensitive phenotype and thus, ASOs targeting *Mogat1* in other tissues does not explain the observed discrepancies. Taken together with previous work, these studies demonstrate the importance of using rigorous and complementary controls when employing ASOs to study intermediary metabolism and insulin sensitivity.

It is now clear that *Mogat1* ASO elicits its effects even in the absence of *Mogat1* expression. In this work and in previous studies, we have used multiple ASO sequences targeting *Mogat1* with similar effects on glucose metabolism and do not anticipate these effects are due to silencing another gene from off-target interactions with RNA. The present results parallel another recent paper that showed that ASOs targeting TTC39B (T39) still elicited metabolic benefits in global T39 KO mice [22]. McCabe and colleagues [22] demonstrated in their model that ASO treatment activated an interferon signaling pathway as indicated by an induction of IFNAR-1 responsive genes (*Oasl1, Ifit1*, and *Ifit2*) that led to adipocyte beiging and caused the mice to lose weight. Mice lacking IFNAR-1 were protected from ASO-induced weight loss and beiging in that study as well. Exactly, how the interferon response is activated by ASO, how this occurs in the KO in the absence of target RNA, and why the scramble control ASO does not provoke the same response is still unclear. Although we did not observe weight loss in our studies, *Mogat1* ASO treatment in WT or MOKO mice stimulated the expression of *Oasl1, Ifit1*, and *Ifit2* in liver and in adipose tissue also induced the expression of *Ucp1* in fat. Moreover, these responses were exaggerated in MOKO mice for reasons yet to be determined. The activation of IFNAR-1 signaling may be consistent with our previous work demonstrating that *Mogat1* ASO exacerbates hepatic inflammation on a diet that induces nonalcoholic steatohepatitis, including components of interferon signaling [21]. While IFNAR-1 signaling activates a multitude of signaling pathways that could plausibly affect glucose metabolism and insulin sensitivity [33], we were unable to prevent the improvements in glucose metabolism by blocking IFNAR-1 activation with an antibody. Thus, IFNAR-1 activation is not the mechanism for improved glucose and insulin tolerance in response to *Mogat1* ASO, which will require further study.

Based on previous work with the *Mogat1* ASO [17,18,30], we were surprised to find that genetic deletion of *Mogat1* in liver did not affect hepatic MGAT activity. Several enzymes exhibit MGAT activity, including *Mogat2* and *Dgat1*, and lack of MGAT activity suppression could be due to compensation from these enzymes [10–13]. *Mogat2* has a higher specific activity than *Mogat1* and in many of the current studies was upregulated by high fat diet feeding. Although we did not find evidence for robust transcriptional activation of these enzymes in the *Mogat1* knockout mice, there could be post-transcriptional mechanisms including post-translational modifications that enhance activity. The lack of effect on hepatic MGAT activity is consistent with previous work showing that global *Mogat1* KO mice do not have deficits in hepatic MGAT activity [34]. However, our previous work has shown that adipocyte-specific KO mice exhibit reduced adipose tissue MGAT activity, which may be consistent with the very low expression of *Mogat2* in adipose tissue [15].

## 5. CONCLUSIONS

Here we provide evidence that *Mogat1* ASO treatment improves whole-body metabolism through *Mogat1*-independent effects. Liver-specific *Mogat1* ablation does not improve insulin resistance in HFD-fed mice, as observed with *Mogat1* ASO treatments [17,18]. Moreover, we show novel evidence that whole-body *Mogat1* deletion leads to insulin resistance, in the context of diet-induced obesity. Finally, we provide preliminary evidence that *Mogat1* ASOs improve glucose intolerance even in global *Mogat1* KO mice. These findings also demonstrate that careful consideration should be given for using ASOs to target gene suppression, including use of genetic knockout models, in metabolic studies that could be affected by these off-target effects.

## ACKNOWLEDGEMENTS

The authors would like to thank Dr. Kyle McCommis at St. Louis University for his insight into off-target effects of ASO treatments, Dr. Eric Yen at the University of Wisconsin for his advice on MGAT biology, and Daniel Ferguson for editing the manuscript during quarantine. We also thank the Washington University School of Medicine Nutrition and Obesity Research Center for continued support in research. Work in the authors’ lab was supported by grants from the NIH (R56 DK111735) and the American Diabetes Association (1-17-IBS-109) to BNF and core laboratories of Washington University School of Medicine (Diabetes Research Center (P30 DK020579), Digestive Diseases Research Cores Center, (P30 DK052574), and the Nutrition Obesity Research Center (P30 DK056341).

